# Coupling metabolic addiction with negative autoregulation to improve strain stability and pathway yield

**DOI:** 10.1101/2020.05.03.075242

**Authors:** Yongkun Lv, Yang Gu, Jingliang Xu, Jingwen Zhou, Peng Xu

## Abstract

Metabolic addiction, an organism that is metabolically addicted with a compound to maintain its growth fitness, is an underexplored area in metabolic engineering. Microbes with heavily engineered pathways or genetic circuits tend to experience metabolic burden leading to degenerated or abortive production phenotype during long-term cultivation or scale-up. A promising solution to combat metabolic instability is to tie up the end-product with an intermediary metabolite that is essential to the growth of the producing host. Here we present a simple strategy to improve both metabolic stability and pathway yield by coupling chemical addiction with negative autoregulatory genetic circuits. Naringenin and lipids compete for the same precursor with inversed pathway yield in oleaginous yeast. Negative autoregulation of the lipogenic pathways, enabled by CRISPRi and fatty acid-inducible promoters, repartitioned malonyl-CoA to favor flavonoid synthesis and increased naringenin production by 74.8%. With flavonoid-sensing hybrid promoters to control leucine synthesis, this flavonoid addiction phenotype confers a selective growth advantage to the naringenin-producing cell. The engineered yeast persisted 90.9% of naringenin titer up to 324 generations. Cells without flavonoid addiction regained growth fitness but lost 94.5% of the naringenin titer after cell passage beyond 300 generations. Metabolic addiction and negative autoregulation may be generalized as basic tools to eliminate metabolic heterogeneity, improve strain stability and pathway yield.

## 1. Introduction

Metabolic heterogeneity has been found to play an essential role in determining cellular performance and pathway efficiency (Xiao, Bowen et al. 2016, Ceroni, Boo et al. 2018, Rugbjerg, Sarup-Lytzen et al. 2018). Traditional bioprocess engineering strategies, including modulating agitation, temperature, pH, dissolved oxygen (DO), dilution rate, and feeding of limiting nutrients, are largely employed nowadays to maintain metabolic homeostasis and improve the titer, yield, and productivity (TYP) of the engineered cell (Qiao, Wasylenko et al. 2017, Xu, Qiao et al. 2017). Fermentation could be viewed as an evolutionary process with productive cells constantly dividing into multiple lineages that are deviating from the parental strains (Xu 2018). In particular, heavily engineered strains with artificial pathways or genetic circuits tend to lose the production phenotype in long-term fermentations due to metabolic burden, enzyme/cofactor imbalance, toxic chemical accumulation, and genetic instability (Wu, Yan et al. 2016). Metabolic burden drives subpopulations of cell to escape the selection pressure, regain growth fitness, propagate and dominate the producing cell line (Lv, Qian et al. 2019). Moreover, this metabolic instability is hard to comply with the high-standard of GMP guidelines in pharmaceutical and biotechnology industry (Madzak 2015).

The decrease and even loss of production has been recently ascribed to nongenetic cell-cell variations or genetic population heterogeneity (Xiao, Bowen et al. 2016, Rugbjerg, Myling-Petersen et al. 2018, Rugbjerg, Sarup-Lytzen et al. 2018). To solve this challenge, we may borrow the concept of “intelligent control” from mechanical or electrical engineering, to encode decision-making feedback functions at the population level and improve the community-level cellular performance. In such a way, the engineered cell could sense the metabolic stress and autonomously adjust cell metabolism to reinforce growth fitness. Indeed, dynamic feedback control has been successfully applied to various organisms to autonomously partition carbon flux and optimize cell metabolism for chemical production in recent years. The development of the metabolator (Fung, Wong et al. 2005), malonyl-CoA inverter gate (Liu, Xiao et al. 2015), malonyl-CoA oscillatory switches (Xu, Li et al. 2014, Xu, Wang et al. 2014, Johnson, Gonzalez-Villanueva et al. 2017, Xu 2019), burden-driven feedback control (Ceroni, Algar et al. 2015, Ceroni, Boo et al. 2018), quorum-sensing based genetic circuits (Gupta, Reizman et al. 2017, Doong, Gupta et al. 2018, Dinh and Prather 2019), antisense RNA-based control (Yang, Lin et al. 2018), and dCas9-CRIRPRi-based circuits (Berckman and Chen 2019, Wu, Chen et al. 2020) have been implemented to relieve metabolic stress and boost cell’s productivity in recent years. To combat cellular heterogeneity, one promising area that is largely underexplored is to consider the mutualistic social interactions and enhance microbial cooperation. We can apply social “punishment-and-reward” rules to eliminate the non-productive cell and incentivize the productive cell (Xiao, Bowen et al. 2016, Lv, Qian et al. 2019). For example, one can link the end-product with cell fitness to selectively enrich the growth of the producing cell-lines (Rugbjerg, Sarup-Lytzen et al. 2018), or use a xenobiotic chemical (i.e., a rare nitrogen/phosphate source or an antibiotics) to confer a competitive growth advantage to the producing cell (Shaw, Lam et al. 2016) and eliminate the growth of the cheater cells during cultivations (Xiao, Bowen et al. 2016). In such a way, we may mitigate the metabolic heterogeneity and stabilize the overproduction phenotype.

The lipogenic acetyl-CoA and malonyl-CoA flux have been repurposed for production of β-carotenoids (Gao, Tong et al. 2017, Larroude, Celinska et al. 2018), polyketides (Markham, Palmer et al. 2018, Liu, Marsafari et al. 2019, Lv, Marsafari et al. 2019, Palmer, Miller et al. 2020), and terpenoids (Jin, Zhang et al. 2019, Zhang, Zhang et al. 2019) in *Yarrowia lipolytica*. However, substantial portion of acetyl-CoA and malonyl-CoA is channeled into lipid synthesis in the engineered strains. These nutraceuticals and lipids compete for the same precursors (acetyl-CoA or malonyl-CoA) with inversed pathway yield in oleaginous yeast, which is a common problem when *Y. lipolytica* is used as chassis to produce acetyl-CoA-derived chemicals (Beopoulos, Nicaud et al. 2011). A large portfolio of engineering work has been dedicated to mitigate lipogenesis or redirect lipid synthesis for heterologous chemical production (Ledesma-Amaro and Nicaud 2016, Markham, Palmer et al. 2018), but the heavily engineered strains, even the chromosomally-integrated cell lines, are often difficult to maintain high performance during long-term cultivations (Roth, Moodley et al. 2009, Xu, Qiao et al. 2017, Wei, Wang et al. 2019). To solve this challenge, we took advantage of the transcriptional activity of metabolite-responsive promoters (Skjoedt, Snoek et al. 2016, D’Ambrosio and Jensen 2017, Wan, Marsafari et al. 2019), aiming to develop an end-product addiction circuit to rewire cell metabolism in *Y. lipolytica*. Coupled with CRISPRi-assisted lipogenic negative autoregulation, the end-product (flavonoid) addiction circuit drives gene expression (leucine synthesis) that is advantageous to productive cell or deleterious to cheater cell. In such a way, we reinforced the fitness of the productive cell and enhanced the overall metabolic performance. Specifically, we have engineered a bilayer dynamic control circuit to autonomously partition metabolic flux and improve strain stability in *Y. lipolytica*. By coupling metabolic addiction with lipogenic negative autoregulation, we improved cell fitness and pathway yield, and the flavonoid-producing phenotype was sustained up to 320 generations. These results highlight the importance of applying dynamic population control and microbial cooperation to improve the community-level metabolic performance.

## 2. Materials and methods

### 2.1 Plasmids and strains

Plasmids pYLXP’-Cre, pYLXP’-Nluc, and pYLXP’-dCas9 were frozen stocks in our lab. The dCas9 contains an SV40 nuclear localization sequence at C-terminal. *Escherichia coli* NEB 5α was used for plasmid construction and maintenance. *Y. lipolytica* Po1f/Δku70 is a derivative of *Y. lipolytica* Po1f (ATCC MYA-2613, MATA ura3-302 leu2-270 xpr2-322 axp2-deltaNU49 XPR2::SUC2) by removing *KU70* gene. *Y. lipolytica* Po1f/Δku70 and NarPro/ASC were used as the chassis.

### 2.2 Molecular biology and genetic cloning

The plasmid pYLXP’-URA3-loxP was constructed in the previous work (Lv, Edwards et al. 2019). The site-directed integration plasmids pΔleu2loxP and pΔxpr2loxP were constructed as follows. The *leu2* upstream and downstream homologous arms leu2-up and leu2-down were amplified from *Y. lipolytica* Po1f genome DNA using primer pairs leu2-up F/leu2-up R and leu2-down F/leu2-down R, respectively. The 600-bp leu2-up fragment was assembled with *Avr*II digested pYLXP’-URA3-loxP using Gibson Assembly, resulting in pΔleu2loxP-up. The 600-bp leu2-down fragment was assembled with *Not*I digested pΔleu2loxP-up using Gibson Assembly, resulting in pΔleu2loxP. The *xpr2* upstream and downstream homologous arms xpr2-up and xpr2-down were amplified from *Y. lipolytica* Po1f genome DNA using primer pairs xpr2-up F/xpr2-up R and xpr2-down F/xpr2-down R, respectively. The 414-bp xpr2-up fragment was assembled with *Avr*II digested pYLXP’-URA3-loxP using Gibson Assembly, resulting in pΔxpr2loxP-up. The 552-bp xpr2-down fragment was assembled with *Not*I digested pΔxpr2loxP-up using Gibson Assembly, resulting in pΔxpr2loxP. pΔleu2loxP and pΔxpr2loxP are YaliBrick plasmids, which can utilize the isocaudarners for fast assembly (Wong, Engel et al. 2017).

Plasmids containing fatty acid inducible promoters were constructed as follows. The 1591-bp *POX2(1591)* promoter was amplified from the genome DNA of *Y. lipolytica* Po1f using primer pair pPOX2(1591) F/pPOX2 (1591) R. The purified PCR product was assembled with *Avr*II/*Xba*I digested pYLXP’ using Gibson Assembly to yield plasmid pPOX2(1591). The 601-bp A1R1 and 438-bp A3-core promoter were amplified from *Y. lipolytica* Po1f genome DNA using primer pairs pPOX2 (1591) F/A1R1 R and A3 F2/pPOX2 (1591) R, respectively. The purified A1R1 and A3-core promoter fragments were assembled with *Avr*II/*Xba*I digested pYLXP’ using Gibson Assembly to yield plasmid pA1R1A3. The 601-bp A1R1-1 and 438-bp A3-core promoter were amplified from *Y. lipolytica* Po1f genome DNA using primer pairs pPOX2 (1591) F/A1R1 R1 and A3 F/pPOX2 (1591) R, respectively. The 601-bp A1R1-2 fragment was amplified from plasmid pA1R1A3 using primer pair A1R1 F2/A1R1 R2. The purified A1R1-1, A1R1-2, and A3-core promoter fragments were assembled with *Avr*II/*Xba*I digested pYLXP’ to yield plasmid p(A1R1)_x2_A3. The resulting plasmids pPOX2(1591), pA1R1A3, and p(A1R1)_x2_A3 will drive the transcription of gRNAs under the control of P_POX2(1591)_, P_A1R1A3_, and P_(A1R1)x2A3_ promoters, respectively.

In order to analyze these inducible promoters, the luciferase encoding gene *Nluc* was used as reporter. *Nluc* was amplified from plasmid pYLXP’-Nluc using primer pair Nluc F/Nluc R. The purified *Nluc* fragment was assembled with *Sna*BI digested pPOX2(1591), pA1R1A3, and p(A1R1)_x2_A3 to yield pPOX2(1591)-Nluc, pA1R1A3-Nluc, and p(A1R1)_x2_A3-Nluc, respectively. The *Avr*II/*Sal*I digested donor plasmids pPOX2(1591)-Nluc, pA1R1A3-Nluc, and p(A1R1)_x2_A3-Nluc were ligated to *Nhe*I/*Sal*I digested destination plasmid pΔleu2loxP to yield pΔleu2loxP-POX2(1591)-Nluc, pΔleu2loxP-A1R1A3-Nluc, and pΔleu2loxP -(A1R1)_x2_A3-Nluc, respectively. Plasmids used in this paper were listed in **Supplementary Table 2**. These plasmids were used to integrate the *Nluc* expressing cassette to the *leu2* site.

Fatty acid synthesis genes *FAS1*, *FAS2*, and *FabD* were used as targeting genes to auto-regulate fatty acid synthesis. To optimize the transcription efficiency, 3 gRNAs targeting different locations were designed for each gene. The gRNA sequences and targeting locations were listed in **Supplementary Table 1**. The 645-bp gRNA_FAS1-1 was synthesized by Genewiz (Frederick, MD). Overlapping sequences were designed at 5’ and 3’ terminals for Gibson Assembly. The 5’ hammer-head ribozyme and 3’ HDV ribozyme sites flanking the gRNA were designed to generate mature gRNAs (Wong, Engel et al. 2017). gRNA_FAS1-1 was assembled with *Xba*I/*Spe*I digested p(A1R1)_x2_A3 using Gibson Assembly to yield plasmid p(A1R1)_x2_A3-FAS1-1. The gRNA upstream xxx-up and downstream xxx-down fragments were amplified using primer pairs gRNA F/gRNA-xxx R and gRNA-xxx F/gRNA R, respectively. Fragments xxx-up and xxx-down were assembled with *Avr*II/*Sal*I digested p(A1R1)_x2_A3 using Gibson Assembly to yield plasmid p(A1R1)_x2_A3-xxx. “xxx” here referred to gRNA. For instance, the upstream fragment FAS1-2-up and downstream fragment FAS1-2-down were amplified from plasmid p(A1R1)_x2_A3-FAS1-1 using primer pairs gRNA F/gRNA-FAS1-2 R and gRNA-FAS1-2 F/gRNA R, respectively (primers used in this study were listed in **Supplementary Table 3**). FAS1-2-up and FAS1-2-down were assembled with *Avr*II/*Sal*I digested p(A1R1)_x2_A3 to yield p(A1R1)_x2_A3-FAS1-2. For all plasmids, the 5’ hammer-head ribozyme and 3’ HDV ribozyme sites flanking the gRNA were designed to generate mature gRNA (Wong, Engel et al. 2017). The *Avr*II/*Sal*I digested donor plasmid pYLXP’-dCas9 was ligated to the *Nhe*I/*Sal*I digested destination plasmid p(A1R1)_x2_A3-xxx to yield p(A1R1)_x2_A3-xxx-dCas9. The *Avr*II/*Sal*I digested donor plasmid p(A1R1)_x2_A3-xxx-dCas9 was subsequently ligated to *Nhe*I/*Sal*I digested destination plasmid pΔleu2loxP to yield pΔleu2loxP-xxx-dCas9. “xxx” here referred to gRNA(s). These plasmids were used to integrate at the *leu2* site. Plasmids used in this paper were listed in **Supplementary Table 2**.

The product addiction plasmids were constructed as follows. The hybrid promoters are composed of FdeR binding sequence *FdeO* and core promoters (Skjoedt, Snoek et al. 2016). *FdeO* was placed at the upstream of the TATA box of *TEF(111)*, *LEU2(78)*, and *GAPDH(88)*, yielding hybrid promoters P_*OTEF(111)*_, P_*OLEU2(78)*_, and P_*OGAPDH(88)*_, respectively. A 25-bp proximal motif was added between *FdeO* and core *TEF* promoter, yielding hybrid promoter P_*OTEF(136)*_ (Hussain, Gambill et al. 2016). These hybrid promoters were synthesized by Genewiz (Frederick, MD), and inserted to pYLXP’2 at *Avr*II/*Xba*I site to replace the original *TEF* promoter, resulting in plasmids pOTEF(111), pOTEF(136), pOLEU2(78), and pOGAPDH(88). Luciferase encoding gene *Nluc* was amplified from pYLXP’-Nluc, and inserted into pOTEF(111), pOTEF(136), pOLEU2(78), and pOGAPDH(88) at *Sna*BI site using Gibson assembly, resulting in pOTEF(111)-Nluc, pOTEF(136)-Nluc, pOLEU2(78)-Nluc, and pOGAPDH(88)-Nluc, respectively. *LEU2* was amplified from pYLXP’ using primer pair LEU2 F/LEU2 R, and inserted into pOTEF(111), pOTEF(136), pOLEU2(78), and pOGAPDH(88) at *Sna*BI site using Gibson Assembly, resulting in pOTEF(111)-LEU2, pOTEF(136)-LEU2, pOLEU2(78)-LEU2, and pOGAPDH(88)-LEU2. *FdeR* containing an SV40 nuclear localization sequence at C-terminal was synthesized by Genewiz (Frederick, MD), and inserted into pYLXP’, pYaliJ1, pYaliL1, pOTEF(111), pOTEF(136), pOLEU2(78), and pOGAPDH(88) at *Sna*BI site using Gibson assembly, resulting in pYLXP’-FdeR, pYaliJ1-FdeR, pYaliL1-FdeR, pOTEF(111)-FdeR, pOTEF(136)-FdeR, pOLEU2(78)-FdeR, and pOGAPDH(88)-FdeR, respectively (Wong, Engel et al. 2017). The recombination of these plasmids was achieved by ligating *Avr*II/*Sal*I digested donor plasmid to *Nhe*I/*Sal*I digested destination plasmid. Plasmids used in this paper were listed in **Supplementary Table 2**.

### 2.3 Site-directed integration

The pΔleu2loxP and pΔxpr2loxP derived plasmids were digested with *Avr*II/*Not*I. The linearized genetic circuits were transformed into *Y. lipolytica* Po1f, NarPro/ACS, or NarPro/ACS_Rep. The uracil drop-out CSM-Ura plate was used to screen transformants. The primer pairs leu2_Inte F/leu2_Inte R and xpr2_Inte F/leu2_Inte R were used to analyze the integration at *leu2* and *xpr2* sites using colony PCR, respectively. The forward primers leu2_Inte F and xpr2_Inte F prime the upstream sequence of the genome, while reverse primer leu2_Inte R primes *URA3* gene in the circuit. The positive colony containing integration at *leu2* site will yield a 1368-bp specific band, while the positive colony containing integration at *xpr2* site will yield a 1556-bp specific band in the colony PCR analysis. The *URA3* marker was rescued by transient expressing Cre recombinase using pYLXP’-Cre, which was subsequently removed by culturing in YPD medium at 30°C for 48 h (Lv, Edwards et al. 2019).

### 2.4 Fatty acid and naringenin inducible promoter analysis

The luciferase encoding gene *Nluc* was used as reporter. The transformant cells were harvested by centrifugation after grown in YPD medium for 24 h. After washing twice using 20 mM PBS, the cells were resuspended in fresh YPD medium. Oleic acid or naringenin was added to concentrations as indicated. The luminescence was recorded using a Cytation 3 microtiter plate reader. The RLU was calculated by integrating the read-out data by time.

### 2.5 Transcriptional assay of fatty acid biosynthesis genes

Reverse transcription - quantitative polymerase chain reaction (qRT-PCR) was used to analysis the transcriptional levels of fatty acid biosynthesis gene. Total RNA was extracted using ToTALLY RNA™ Kit (Ambion, Inc.) at 48 h. DNA was removed using TURBO DNA-*free*™ Kit (Invitrogen, Vilnius, Lithuania). Reverse transcription was carried out using an iScript™ cDNA Synthesis kit (Bio-Rad, Hercules, CA). Real time PCR was carried out using an iTaq™ Universal SYBR^®^ Green Supermix (Bio-Rad, Hercules, CA) using the Bio-Rad iCycler iQ Real-Time PCR Detection System (Bio-Rad, Hercules, CA). The fluorescence results were analyzed using Real-time PCR miner. Actin encoding gene *ACT1* (GRYC ID: YALI0D08272g) was used as reference gene (Tai and Stephanopoulos 2013). Relative changes in gene transcriptional level were calculated using ΔΔC_q_ method (Livak and Schmittgen 2001). Primers used for qRT-PCR were listed in **Supplementary Table 3**.

### 2.6 Month-long fermentation test of the naringenin-producing cell line

The month-long fermentation testing was implemented by serial passaging in shaking flasks (Xiao, Bowen et al. 2016, Rugbjerg, Sarup-Lytzen et al. 2018). Inoculum seedling culture aging from 0 hour to 720 hours were individually harvested and stocked in −80°C freezer before the inoculum was used as seed culture. Then the individually harvested seeds were revived on solid agar minimal media (CSM-Leu) before inoculation into 3 ml liquid media. To avoid cell density variations, the liquid seed with an initial OD 0.1 (normalized by the individual culture) was inoculated into 25 ml fresh CSM-Leu medium in 250 ml shaking flask. Samples were taken and stored in −80°C for flavonoid analysis. To analyze the naringenin productivity, 1 ml of the −80°C frozen culture harvested at 120 h was used to analyze naringenin titer with HPLC (Lv, Marsafari et al. 2019).

## 3. Results and discussion

### 3.1 Construction of a fatty acid-driven inverter gate

We previously engineered the flavonoid biosynthetic pathway in *Y. lipolytica* by overexpressing the acetyl-CoA carboxylase and supplementing media with the expensive FAS inhibitor (1 mg/L cerulenin) (Lv, Edwards et al. 2019). However, the engineered cell still contained more than 35% (w/w) of oil (primarily neutral lipids, triacyl glycerides) (Beopoulos, Nicaud et al. 2011), indicating a substantial portion of malonyl-CoA was used for lipid synthesis. Consistent with this study, mitigating lipogenesis competition has been reported as effective strategies to improve malonyl-CoA-derived end product synthesis in oleaginous yeast (Wu, Yu et al. 2014, Yang, Lin et al. 2015). Since lipogenesis is essential to membrane synthesis and cellular function, it is important to balance the flux between lipogenesis and heterologous product synthesis.

As dynamic control has emerged as effective strategy to regulate metabolic flux (Xu 2018), we sought to build a fatty acid-driven inverter device to autonomously tune down lipids synthesis when there is sufficient fatty acids in the cell. To autonomously tune down fatty acid synthesis, we engineered a fatty acid inducible promoter in combination with the CRISPRi elements to repress lipogenic pathways (Fig. 1). Previous report has validated the essential molecular components of a fatty acid inducible promoter *POX2*, which is composed of upstream activating sequences (A1, A2, and A3), regulatory sequences (R1 and R2), and the core promoter sequence (**Fig. 2a**) (Hussain, Wheeldon et al. 2017). Both the intact *POX2* promoter and its derivatives have been reported to be inducible by fatty acids (Hussain, Gambill et al. 2016, Hussain, Wheeldon et al. 2017). We tested a number of genetic configurations of the *POX2* promoter to improve the dynamic output range and operational range (Fig. 2). When induced by 2% (v/v) oleic acids, the hybrid promoter *(A1R1)_x2_A3* exhibited the highest gene expression fold change with an engineered luciferase (encoded by *Nluc*) as the reporter (**Fig. 2b**). Moreover, the hybrid promoter *(A1R1)_x2_A3* was characterized to respond to oleic acids ranging from 0.1% (v/v) to 5.0% (v/v) (**Fig. 2c**). Considering the dynamic output and the operational range of the hybrid promoter, the hybrid promoter P_*(A1R1)x2A3*_ was chosen to drive the expression of guide RNAs (gRNAs) that were specifically designed to target the lipogenic pathways. The 5’ hammer-head ribozyme (5’ HHR) and 3’ HDV ribozyme sites flanking the gRNAs were used to generate mature gRNA (Wong, Engel et al. 2017). Due to plasmid instability in *Y. lipolytica*, we integrated the inverter device at the *leu2* loci by using site-specific integration plasmid pΔleu2loxP (**Supplementary Fig. 1a, 1b and 1c**) (Le Dall, Nicaud et al. 1994). The integration efficiency was determined to be 67.7% (21/31) in *KU70* deficient strain (**Supplementary Fig. 1b**). By transiently expressing the Cre recombinase, we were able to efficiently cure the *URA3* marker with 100% (23/23) efficiency (**Supplementary Fig. 1c**) (Lv, Edwards et al. 2019). This site-specific gene integration enable us to develop a stable inverter device to autonomously control lipid synthesis, and integration of the inverter gate at *leu2* loci will further block the native leucine biosynthetic pathway, rendering us the opportunity to link leucine auxotroph (growth fitness) with flavonoid synthesis (which will be described in section 2.4).

**Fig. 1.**
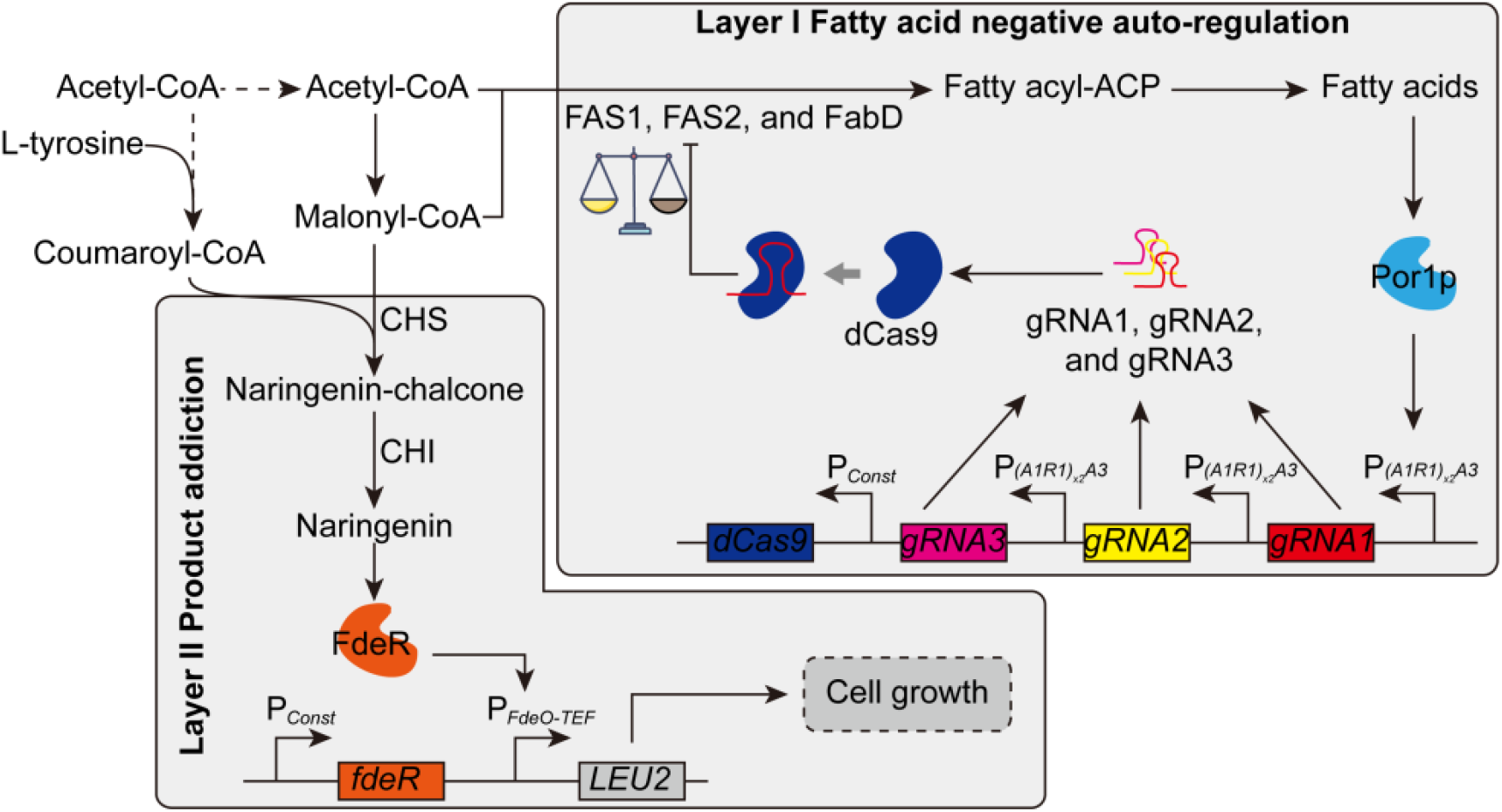
Demonstration of integrating fatty acid negative auto-regulation with product-addiction. Layer I: CRISPRi based fatty acid negative auto-regulation. The fatty acid inducible promoter P_*(A1R1)x2A3*_ was used to control the transcription of gRNAs, which were used to guide dCas9 to *FAS1*, *FAS2*, and *FabD*. Layer II: End-product addiction. P_*FdeO-TEF*_ is a hybrid promoter, which is composed of FdeR binding site *FdeO* and the *TEF* core promoter. P_*Const*_ refers to the constitutive promoter. In the presence of naringenin, FdeR bands *FdeO* site and activates P_*FdeO-TEF*_. The expression of *LEU2* gene confers the leucine-auxotrophic host cell growth in the leucine drop-out medium.

**Fig. 2.**
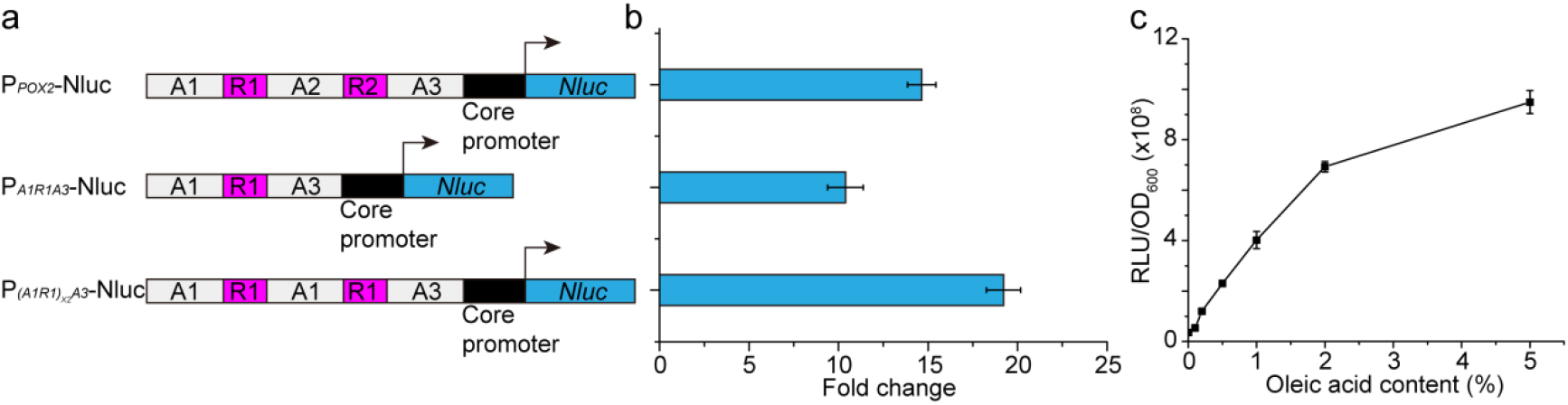
Analysis of fatty acid inducible promoters. **a** Structures of the fatty acid inducible promoters. **b** Dynamic output range of fatty acid inducible promoters. The fold change was calculated by dividing the RLU/OD_600_ with 2% (v/v) oleic acid by the RLU/OD_600_ without oleic acid. **c** Operational range of hybrid promoter *(A1R1)_x2_A3*. The oleic acid content was the volume content.

### 3.2 CRISPR interference to inhibit fatty acid synthesis

Yeast uses polypeptide fatty acid synthetases Fas1p and Fas2p (encoded by *FAS1* and *FAS2*) to assemble acetyl-CoAs and malonyl-CoAs into long-chain saturated fatty acids (Schweizer, Roberts et al. 1986). The *Y. lipolytica* homologue of malonyl-CoA-ACP [acyl-carrier protein] transacylase (encoded by *FabD*, GRYC ID: YALI0E18590g) was also reported to play a major role in fatty acid synthesis (Schneider, Brors et al. 1997). For these reasons, *FAS1*, *FAS2*, and *FabD* were selected as endogenous CRISPRi targets to repress fatty acid synthesis. In order to achieve gradient repression levels, we tested 3 gRNAs targeting different locations for each gene (**Supplementary Table 1**). The hybrid promoter P_*(A1R1)x2A3*_ was used to drive the transcription of gRNAs. The mature gRNAs, which were used to guide dCas9 to the target locations, were produced by endogenous processing of the flanking 5’ hammer-head ribozyme and 3’ HDV ribozyme sites after transcription (**Fig. 3a-c**).

**Fig. 3.**
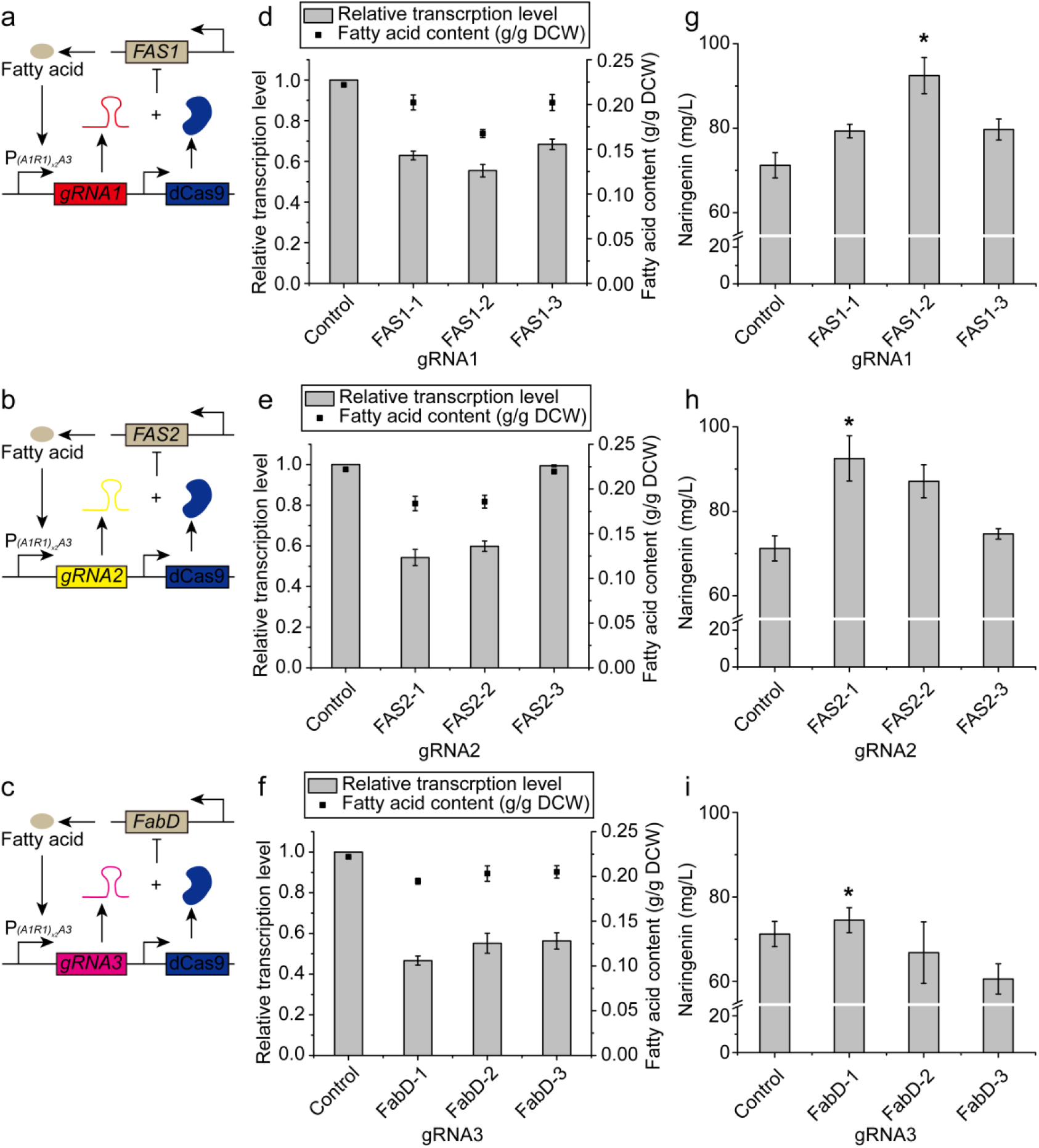
Effects of different gRNAs on relative transcription level, fatty acid content, and naringenin production. **a** - **c** Mechanism of gRNA1, gRNA2, and gRNA3 guided fatty acid synthesis auto-regulation. **d** - **f** Effects of different gRNA1, gRNA2, and gRNA3 on relative transcription level and fatty acid content. The relative transcription level of *FAS1*, *FAS2*, and *FabD* was normalized by the transcription level of *FAS1*, *FAS2*, and *FabD* without auto-regulation (the control). **g** - **i** Effects of repressing *FAS1*, *FAS2*, and *FabD* using different gRNA1, gRNA2, and gRNA3 on naringenin titer. Strain NarPro/ASC without auto-regulation circuit was used as control. gRNA1 refers to gRNA FAS1-1, FAS1-2, or FAS1-3. gRNA2 refers to gRNA FAS2-1, FAS2-2, or FAS2-3. gRNA3 refers to FabD-1, FabD-2, or FabD-3.

CRISPRi experiments showed that different gRNAs resulted in differential transcriptional repression for each gene (**Fig. 3d-f**). The gRNAs FAS1-2, FAS2-1, and FabD-1 showed the highest transcriptional repression efficiency, resulting in relative transcription levels of 0.55, 0.54, and 0.47, respectively, as quantified by qRT-PCR. Consistent with these transcriptional data, the gRNAs FAS1-2, FAS2-1, and FabD-1 guided dCas9 decreased fatty acid content by 24.5%, 17.2%, and 12.3% respectively (**Fig. 3d-f**). gRNAs targeting *FabD* was less efficient in decreasing fatty acid accumulation, possibly due to the fact that FabD activity was compensated by the pentafunctional Fas1p (Schweizer, Roberts et al. 1986, Kottig, Rottner et al. 1991). Both gRNAs FAS1-2 and FAS2-1 were found to increase naringenin titer by 29.9%, while the gRNA FabD-1 did not have obvious effect on naringenin titer (**Fig. 3g-i**). These results indicated that naringenin production was improved by dCas9 interfering with fatty acid synthesis when the gRNAs were controlled by the fatty acid inducible promoter.

### 3.3 Fatty acid synthesis negative autoregulation by multiplexed gRNAs-CRISPRi tuning

To further divert lipogenic flux toward the flavonoid pathway, we sought to use gRNAs in combination to target *FAS1*, *FAS2*, and *FabD* and test whether the multiplexed gRNAs could improve naringenin production (**Fig. 4a**). The results showed that combinatorially repressing *FAS1-2* and *FAS2-1* decreased fatty acid dramatically comparing with repressing single gene, while the duplex gRNAs targeting *FAS1-FabD* or *FAS2*-*FabD* did not result in significant fatty acid reduction comparing with solo repression of *FAS1* or *FAS2* (**Fig. 4b**). Duplex gRNAs targeting *FAS1*-*FAS2* increased naringenin titer by 52.8%, and triplex gRNAs targeting *FAS1*-*FAS2*-*FabD* increased naringenin titer by 56.5% (**Fig. 4b**). We further confirmed that naringenin titer was negatively correlated with fatty acid content (R^2^ = 0.98) (**Fig. 4c**). These results indicated that naringenin production was improved with the multiplexed gRNAs targeting the fatty acid pathway. Specifically, the strain NarPro/ASC_Rep produced 111.4 mg/L naringenin (**Fig. 4b**), when triplex gRNAs (FAS1-2-FAS2-1-FabD-1) were used to repress the fatty acid pathway.

**Fig. 4.**
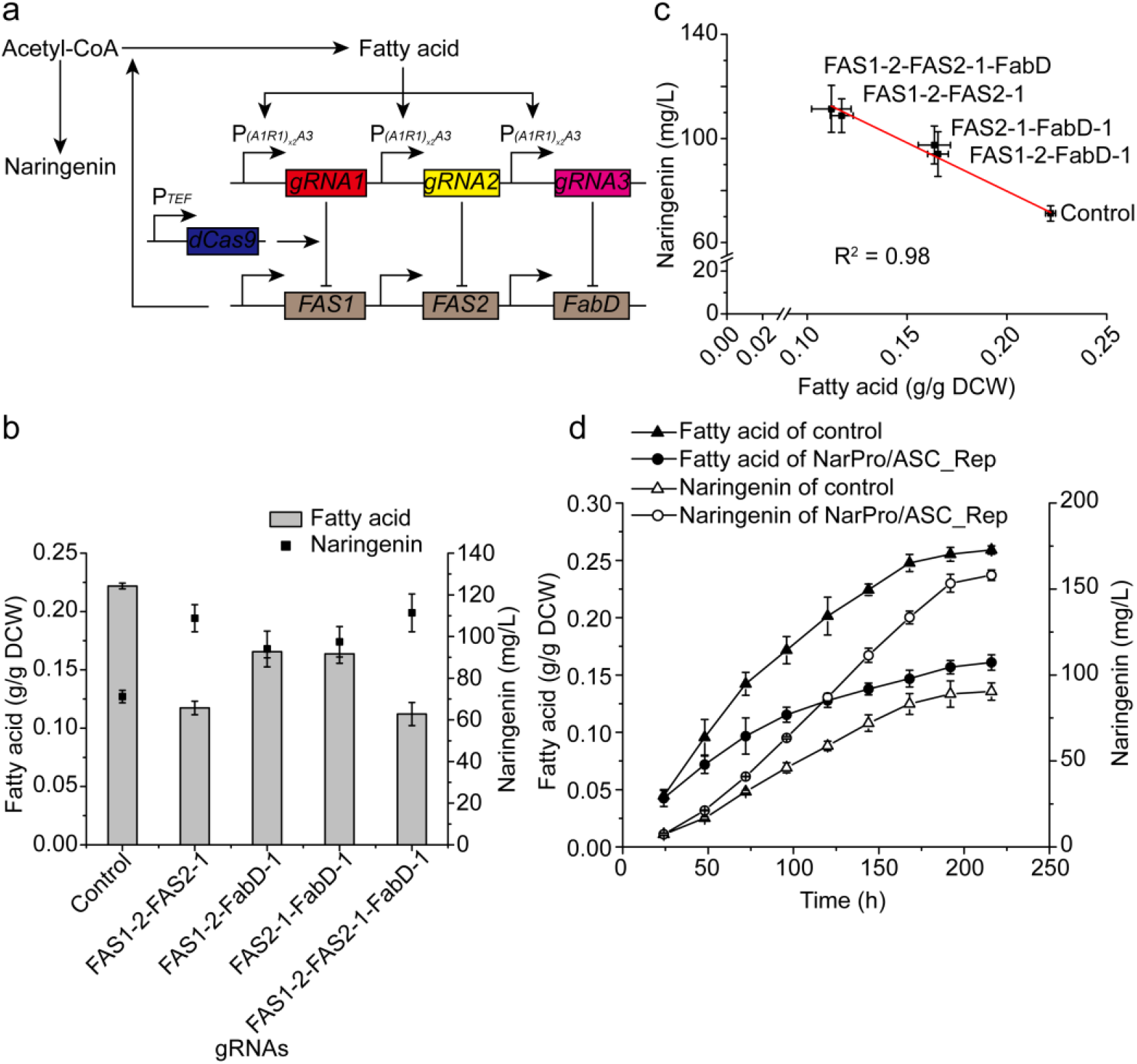
Combinatory repression analysis and time course of fatty acid and naringenin production. **a** Mechanism of auto-regulating fatty acid synthesis by combinatory repressing *FAS1*, *FAS2*, and/or FabD. **b** Effects of combinatory repressing *FAS1*, *FAS2*, and/or FabD on fatty acid accumulation and naringenin titer. **c** Analysis of the tradeoff between fatty acid and naringenin synthesis. The red line refers to the linear fit to the mean values. **c** Time course of fatty acid accumulation and naringenin production. Strain NarPro/ASC, which is identical to NarPro/ASC_Rep but without the negative auto-regulation circuit, was used as control.

We further analyzed the time course of fatty acid and naringenin production using 250-mL shaking flask in fed-batch fermentation. Interestingly, fatty acid accumulation was reduced further in NarPro/ASC_Rep than the control strain. At the end of the fermentation, fatty acid in NarPro/ASC_Rep decreased by 37.9% (**Fig. 4d**). Naringenin production rate was increased substantially in the NarPro/ASC_Rep strain. At the end of the fermentation, NarPro/ASC_Rep produced 158.0 mg/L naringenin, which was 74.8% higher than that of the control (**Fig. 4d**). The inverse correlation between naringenin and fatty acid confirmed that the triplex FAS1-2-FAS2-1-FabD-1 gRNAs effectively guided dCas9 to divert the lipogenic carbons toward the naringenin pathway. In summary, the fatty acid-driven negative auto-regulation inverter gate provided a promising solution to mitigate precursor flux competition from lipogenesis in *Y. lipolytica*.

### 3.4 Construction of naringenin inducible promoters

Despite the improvement of naringenin production by mitigating fatty acid synthesis, we constantly observed that the engineered yeast gradually lost the production phenotype after several generations of cultivation. To solve this problem, we attempted to engineer an end-product (naringenin) addiction circuit, which will link cell growth fitness with naringenin production. This competitive growth advantage will encourage the proliferation of the high naringenin-producing populations, while suppress or eliminate the growth of the low naringenin-producing populations during long-term cultivation. To link cell fitness with the end-product, we sought to place an essential gene under the control of naringenin inducible circuit. We firstly ruled out the antibiotic resistant genes, because the antimicrobial property of both antibiotics and naringenin may make the addiction circuit difficult to validate. Because the *LEU2* gene is a ready to use (the chassis is leucine-auxotrophic) and reliable selective marker, we firstly tried this non-conditionally essential gene. Cultivated with the leucine drop-out synthetic medium, genetic variants with high naringenin titer will produce more leucine and outcompete the growth of the low naringenin-producing strain. As a result, the high naringenin-producing strains will gradually dominate the cell populations and low naringenin-producing cells will be suppressed (Layer II in **Fig. 1**). To build the naringenin-inducible genetic circuits, we firstly sought to use the well-characterized transcriptional activator FdeR and its cognate DNA binding site *FdeO* (Skjoedt, Snoek et al. 2016) to control the expression of a reporter gene. A panel of synthetic promoters P_*O-TEF(111)*_, P_*O-TEF(136)*_, P_*O-LEU2(78)*_, and P_*O-GAPDH(88)*_ were constructed by fusing *FdeO* with the core promoters to drive the expression of the *Nluc* luciferase (**Fig. 5a**). The nuclear localization signal SV40 was fused at the C-terminal of FdeR to facilitate nuclear transportation. With naringenin as the effector molecule, FdeR will bind to *FdeO* site and activate the transcription of the reporter gene (**Fig. 5b**) (Siedler, Stahlhut et al. 2014). Experimental results demonstrated that all our constructed promoters were inducible by naringenin with an operational range from 0 mg/L to 50 mg/L (**Fig. 5c**). The induction reached saturation when naringenin concentration was beyond 100 mg/L. These naringenin-inducible promoters will be used to control the expression of essential gene *LEU2* that dictates cell fitness.

**Fig. 5.**
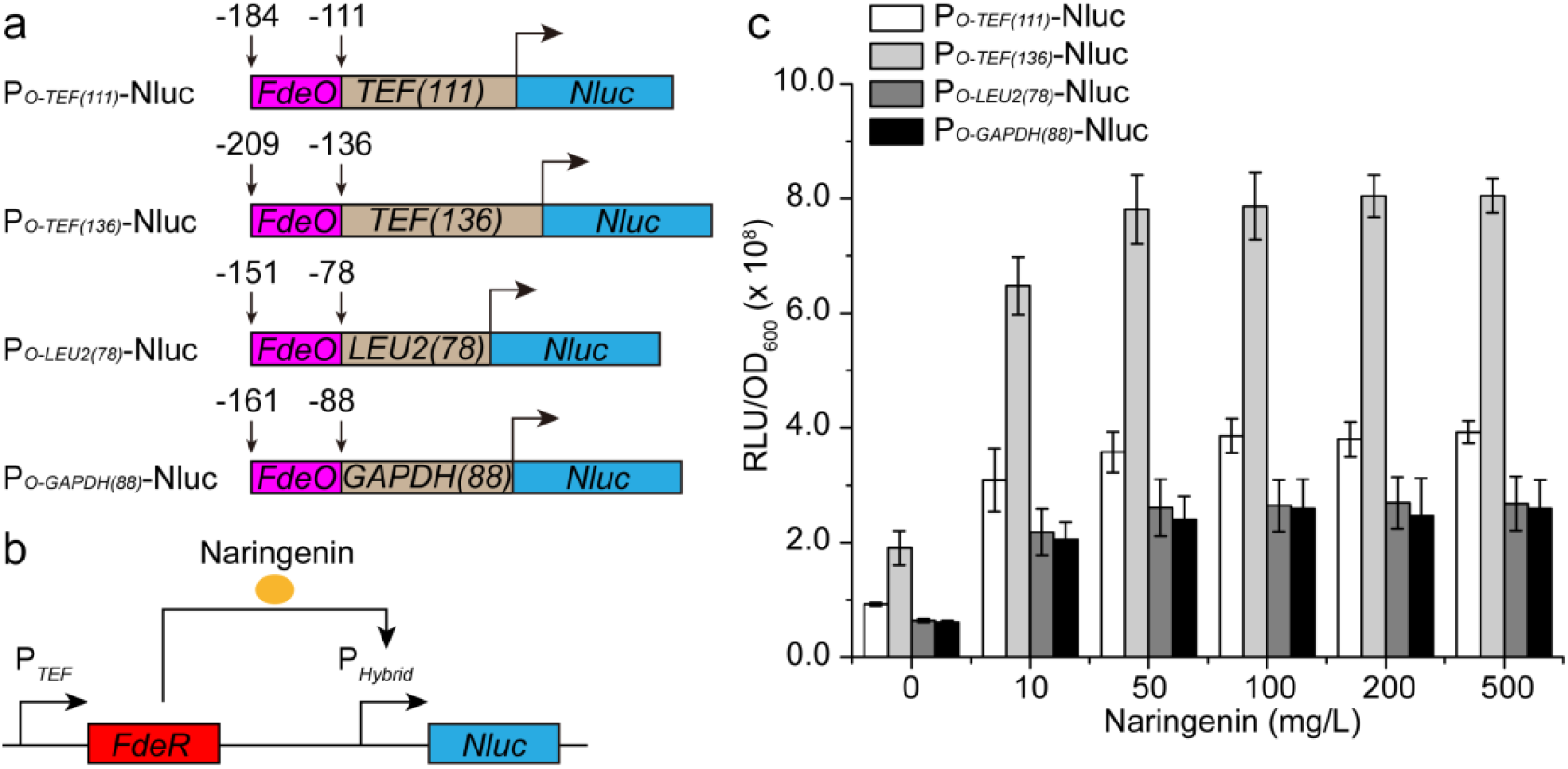
Naringenin inducible promoter analysis. **a** Structures of the hybrid promoters. The hybrid promoters *OTEF(111)*, *OTEF(136)*, *OLEU2(78)*, and *OGAPDH(88)* were composed of FdeR binding site *FdeO* and the core promoters of *TEF(111)*, *TEF(136)*, *LEU2(78)*, and *GAPDH(88)* respectively. **b** Mechanism of the naringenin inducible transcription. P_*Hybrid*_ refers to the hybrid promoters of P_*O-TEF(111)*_, P_*O-TEF(136)*_, P_*O-LEU2(78)*_, and P_*O-GAPDH(88)*_. *Nluc* was used as the reporter gene. **c** The result of induction of the hybrid promoters. The luminescence was recorded using Cytation 3 microtiter plate reader. The RLU was calculated by integrating the read-out data by time.

### 3.5 Linking flavonoid production with cell growth fitness

One important strategy in metabolic engineering is to engineer competitive growth advantage that links cell growth with product formation. To enable naringenin inducible growth, we placed the leucine biosynthesis gene *LEU2* under the control of these naringenin-inducible promoters (**Supplementary Fig. 2a-d**). Upon transformation into the *leu2*-deficient *Y. lipolytica* Po1f chassis, we obtained comparable number of colonies on the plates with naringenin (CSM-Leu+Naringenin) or without naringenin (CSM-Leu) (**Supplementary Fig. 2e-l**), indicating that cell growth is independent of naringenin. We speculate this might be arising from the basal activity of *FdeR*, which was excessively expressed from the strong and constitutive *TEF* promoter. To overcome this limitation, we next sought to express FdeR with weak promoters *POX4* and *IDP2* (Wong, Engel et al. 2017). Indeed, we observed naringenin-inducible growth: FdeR expression from the weak promoters (P_*POX4*_ and P_*IDP2*_) effectively activated the leucine circuits (P_*O-TEF(136)*_-*LEU2*) and supported a distinguishable growth phenotype in the presence of naringenin (**Fig. 6d, e, g, and h**). However, when leucine circuits P_*O*-*TEF(111)*_-*LEU2*, P_*O*-*LEU2(78)*_-*LEU2*, and P_*O*-*GAPDH(88)*_-*LEU2* were tested, none of them could grow on CSM-Leu or CSM-Leu+naringenin plates, indicating an intricate interaction between FdeR and these synthetic promoters. Interestingly, FdeR expression from the naringenin-inducible promoter (P_*O-TEF(136)*_) also conferred naringenin-inducible growth (**Fig. 6f and i**), this possibly created a positive feedforward loop that may amplify the sensor input-output relationship. We next confirmed the naringenin-inducible growth in liquid media. Indeed, *Y. lipolytica* strains carrying the naringenin inducible circuits grew faster in CSM-Leu+naringenin medium than those in CSM-Leu medium (**Fig. 6j and l**), indicating the naringenin inducible circuits conferred a competitive growth fitness for the engineered cell, albeit flavonoids have been reported as antimicrobial agents in plant signaling. These results highlighted the importance of adjusting promoter strength to achieve the desirable end-product addiction phenotype. For example, excess expression of FdeR instead led to naringenin-independent expression of *LEU2*, causing indistinguishable cell fitness. While too less FdeR is not sufficient to activate the *LEU2* circuits and support cell growth. As for the *LEU2* expression, it is important to select the naringenin-inducible promoter with broad operational range as well as minimal leaky expression.

**Fig. 6.**
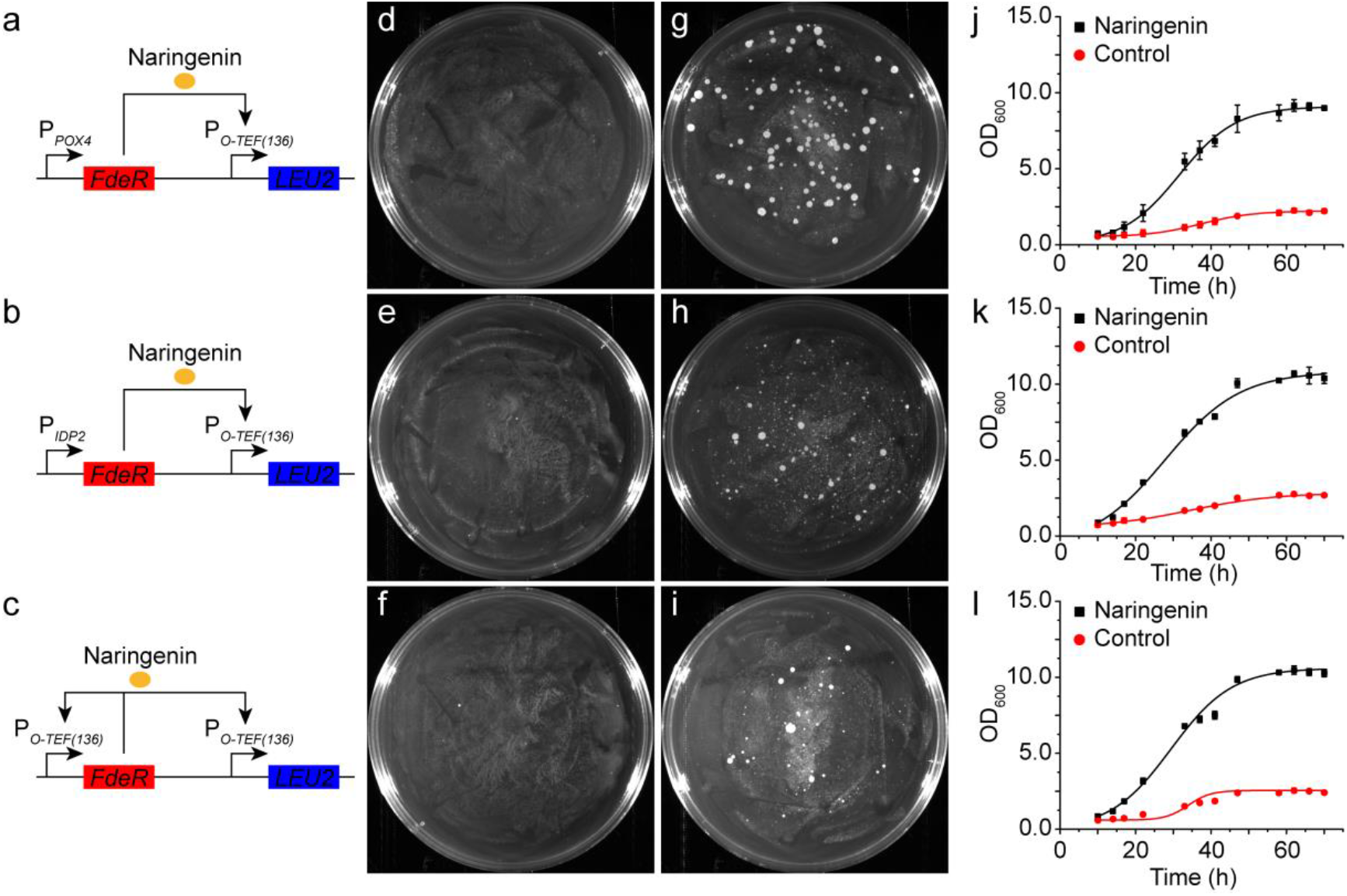
Analysis of naringenin addiction circuits in *Y. lipolytica* Po1f. **a-c** Mechanism of addiction circuits of P_*O-TEF(136)*_-LEU2-P_*POX4*_-FdeR, P_*O-TEF(136)*_-LEU2-P_*IDP2*_-FdeR, and P_*O-TEF(136)*_-LEU2-P_*O-TEF(136)*_-FdeR. **d-f** Growth of *Y. lipolytica* Po1f transformants with P_*O-TEF(136)*_-LEU2-P_*POX4*_-FdeR, P_*O-TEF(136)*_-LEU2-P_*IDP2*_-FdeR, and P_*O-TEF(136)*_-LEU2-P_*O-TEF(136)*_-FdeR circuits on CSM-Leu plates. Pictures were taken 3 days after transformation. **g-i** Growth of *Y. lipolytica* Po1f transformants with P_*O-TEF(136)*_-LEU2-P_*POX4*_-FdeR, P_*O-TEF(136)*_-LEU2-P_*IDP2*_-FdeR, and P_*O-TEF(136)*_-LEU2-P_*O-TEF(136)*_-FdeR circuits on CSM-Leu+Naringenin plates. Pictures were taken 3 days after transformation. **j-l** Growth curves of *Y. lipolytica* Po1f equipped with P_*O-TEF(136)*_-LEU2-P_*POX4*_-FdeR, P_*O-TEF(136)*_-LEU2-P_*IDP2*_-FdeR, and P_*O-TEF(136)*_-LEU2-P_*O-TEF(136)*_-FdeR circuits in liquid CSM-Leu medium with or without naringenin. The control set was in CSM-Leu liquid medium without naringenin. The experimental set was in CSM-Leu liquid medium with 5 mg/L naringenin.

To link cell fitness with naringenin production, we next transformed the naringenin-inducible genetic circuit into the naringenin producing strain NarPro/ASC_Rep with fatty acid negative autoregulation. The fatty acid negative autoregulatory circuits were integrated at the *leu2* genomic loci. To validate naringenin-inducible growth, we observed that strains carrying the naringenin-addicting circuits grew normally on leucine drop-out plates (**Supplementary Fig. 3e-h**). However, the strains grew poorly or did not grow at all when the transcriptional activator FdeR was removed from the system (**Supplementary Fig. 3i-l**), indicating the tight control of the *LEU2*-expressing fitness circuits by FdeR. In the liquid media, strains carrying the addiction circuits enabled the cell reach 3-4 fold higher cell density (OD_600_) with 5 mg/L naringenin than the same strain without naringenin (**Fig. 6 j, k, l**).

### 3.6 Stabilizing naringenin-production phenotype by integrating metabolic addiction with negative autoregulatory circuits

We next coupled the metabolic addiction with negative autoregulation, and validated the long-term robustness of the engineered strains with a month-long fermentation experiment. Three naringenin addiction circuits (P_*O-TEF(136)*_-LEU2-P_*POX4*_-FdeR, P_*O-TEF(136)*_-LEU2-P_*IDP2*_-FdeR, and P_*O-TEF(136)*_-LEU2-P_*O-TEF(136)*_-FdeR) were introduced to the naringenin producing strain with fatty acid negative autoregulation (NarPro/ASC_Rep chassis), which drives leucine synthesis in a naringenin-dependent manner. By incorporating the naringenin-addiction circuits, we locked the naringenin production phenotype with cell growth fitness. The NarPro/ASC_Rep chassis with blank pYLXP’ plasmid was used as control. These engineered strains were undergone serial passage in paralleled shaking flasks experiments (Rugbjerg, Sarup-Lytzen et al. 2018). Cells harvested from 0 to 720 hours (1 month, about 360 generations) were individually stocked and used as seed culture to inoculate the leucine dropout media (CSM-Leu). We confirmed that the strains carrying the addiction circuits sustained the naringenin production phenotype compared with the control strains (**Fig. 7a, 7b, 7c** and **7d**), albeit the control strain (without naringenin-addiction circuit) regained growth fitness and reached relatively higher cell density (Fig. 7a). For example, the control stain and all the addiction strains produced similar amount of naringenin (80-90 mg/L) for the inoculum aging from 0-180 generations (2 hour per generation, which is about 360 hours or 15 days). After 200 generations, naringenin production declined rapidly in the control strain, compared with the strains with addiction circuits (Fig. 7b,7c and 7d). For the inoculum at 648 hours (which is about 324 generations), all the three strains carrying the addiction circuits maintained 90.9% of naringenin production compared with the inoculum at 0 hour. On the contrary, the control strain (without the naringenin addiction circuits) almost lost all the naringenin, only produced less than 5.5% of naringenin compared with the fresh inoculum (0 hour). By linking leucine synthesis with flavonoid production, our coupling strategies effectively force the cell to maintain a selective growth advantage to enrich the flavonoid-producing populations, but suppress the mutant cells that produce less naringenin. These results demonstrated that the end-product addiction circuits are invaluable tools to maintain community-level strain stability, which is a critical index to stabilize recombinant metabolite production and prevent genetic degeneration during bioprocess scale-up and long-term fermentation.

**Fig. 7.**
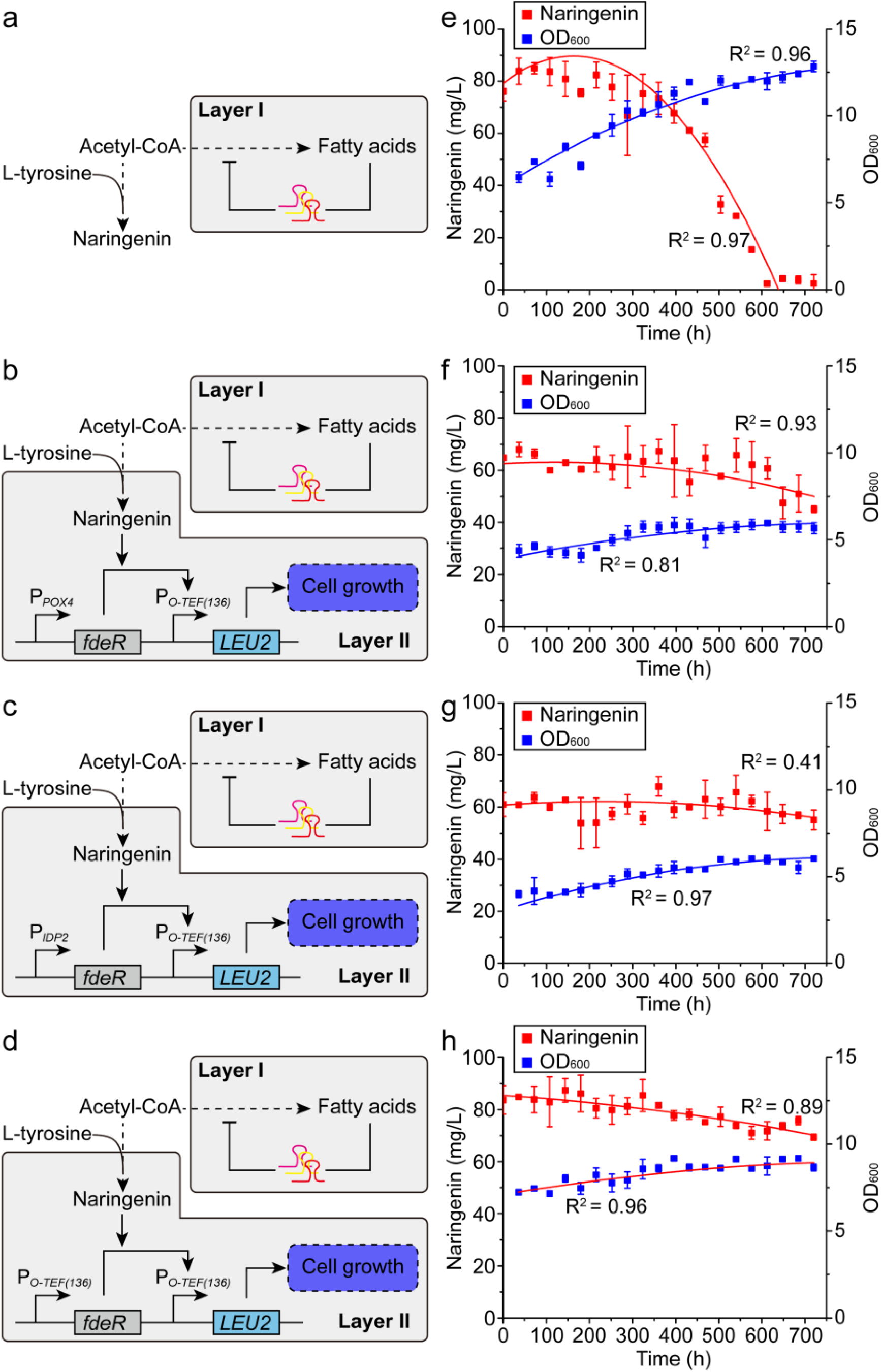
Stability analysis of naringenin producing strains equipped with or without naringenin-addiction circuits. The metabolic addiction were coupled with fatty acid negative autoregulation by equipping naringenin addiction circuits (P_*O-TEF(136)*_-LEU2-P_*POX4*_-FdeR, P_*O-TEF(136)*_-LEU2-P_*IDP2*_-FdeR, and P_*O-TEF(136)*_-LEU2-P_*O-TEF(136)*_-FdeR) to the NarPro/ACS_Rep chassis, which is a naringenin producing strain with fatty acid negative autoregulatory circuit. **a.** Stability analysis of the control strain (NarPro/ACS_Rep chassis with blank pYLXP’ plasmid). **b.** Stability analysis of NarPro/ACS_Rep equipped with P_*O-TEF(136)*_-LEU2-P_*POX4*_-FdeR. **c.** Stability analysis of NarPro/ACS_Rep equipped with P_*O-TEF(136)*_-LEU2-P_*IDP2*_-FdeR. **d.** Stability analysis of NarPro/ACS_Rep equipped with P_*O-TEF(136)*_-LEU2-P_*O-TEF(136)*_-FdeR. Long-term fermentation was carried out in CMS-leu synthetic drop out medium. The strains were passaged at exponential phase (every 36 h). Before every passaging, OD_600_ was measured and samples were taken for frozen stocks. To analyze the naringenin production, the frozen stocks of each sample were re-inoculated into CSM-leu synthetic drop out medium (4% (v/v)). Naringenin titer was measured at 120 h. The lines were the polynomial fits to the means.

Nonetheless, the overall naringenin production is decreased after we combine the fatty acid negative autoregulatory circuit and the flavonoid addiction circuit (Fig. 7a and 7b). The decrease in naringin production is possibly due to the metabolic overloading or burden effect (Wu, Yan et al. 2016) due to the expression of multiple transcriptional regulators (CRISPRi, FdeR and Por1p, and three gRNAs). From an evolutionary perspective, there is a tradeoff between metabolic stability and pathway yield. Metabolites (i.e. leucine or naringenin in this study) cross-feeding could largely elicit metabolic heterogeneity (Evans, Kempes et al. 2020), for example, leucine secreted from the high naringenin-producing strain may be assimilated by the low naringenin-producing strain, therefore causing a faulted cross-talk between the sub-populations of the engineered cell. To restrict leucine cross-feeding, we may need to further delete the gene encoding leucine permease (Bap2 homologs) to sequestrate leucine and reinforce the naringenin production phenotype. Single cell imaging and genetic analysis will be critical to help us understand the source of this metabolic heterogeneity and propose novel genetic or process engineering solutions to inhibit such metabolic heterogeneity (Hartline, Mannan et al. 2020).

## 4. Conclusions

Metabolic heterogeneity has become a major issue that compromises the cellular performance and pathway yield. To overcome this limitation, it is important to rewire cellular logics to maintain metabolic homeostasis and improve the community-level cell performance. Social reward-punishment rules to incentivize the production cell and punish the nonproduction cell may be applied to combat this fitness loss. Such rules could be biologically implemented by conferring a selective growth advantage to the production cell, in such a way, the population of the production cell will be enriched and therefor the production phenotype might be sustained. Most of the reported strategies have focused on nongenetic cell-to-cell variations to confer competitive fitness. Genetic underpinnings that are associated with metabolic heterogeneity remain a challenging area. Single-cell analysis and microfluidic-integrated optogenetic tools might be a promising area to solve these conundrums. In this work, we have engineered lipogenic negative autoregulation with metabolite addiction to redistribute carbon flux and improve strain stability in *Y. lipolytca*. The flavonoid-producing phenotype was sustained for up to 320 generations. These results highlight the importance of applying dynamic population control and microbial cooperation to improve the community-level metabolic performance, which might be useful to combat metabolic heterogeneity and stabilize overproduction phenotype for long-term cultivation in industrial biotechnology settings.

## Acknowledgement

This work was supported by the Cellular & Biochem Engineering Program of the National Science Foundation under grant no.1805139 and the Bill and Melinda Gates Foundation (OPP1188443). YL would like to thank the China Scholarship Council, China Postdoctoral Science Foundation (Grant No. 2019M662535), and Science and Technology Project of Henan Province (Grant No. 202102310019) for funding support.

## Author contributions

PX and YL conceived the topic and designed the study. YL performed genetic engineering and fermentation experiments with output from YG. YL and PX wrote the manuscript. YL is a joint scientist supervised by JWZ, JLX and PX.

