## Supplemental Tables and Figures for "Coupling metabolic addiction with negative autoregulation to improve strain stability and pathway yield"

### Supplementary figures

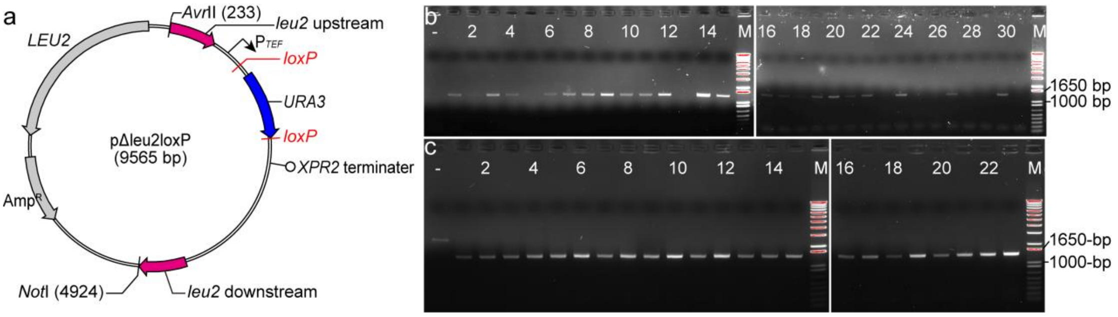


**Supplementary Fig. 1** Site-directed integration at *leu2* loci and *URA3* marker curation. **a** Structure of plasmid pΔleu2loxP. *leu2* upstream and *leu2* downstream refer to the up and down homologous sequence of *leu2* loci. The *loxP* sites were used for *URA3* marker curation. **b** Colony PCR analysis of site-directed integration at *leu2* site. The colony PCR was carried out using primer pair leu2_Inte F/leu2_Inte R. The positive colony will yield a 1368-bp band, while the negative colony will not yield any specific band. **c** Colony PCR analysis of *URA3* marker curation. The colony PCR was carried out using primer pair 26srDNA2s F/XPR2_Seq. The positive colony will yield a 1533-bp band, while the negative colony will yield a 2432-bp band.

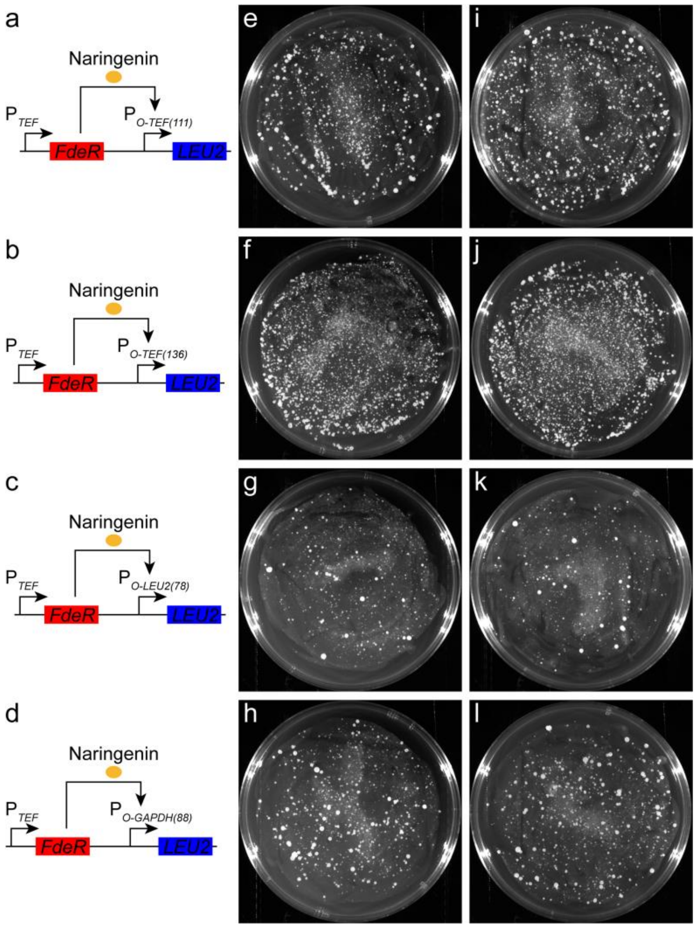


**Supplementary Fig. 2** Analysis of naringenin addiction circuits on CSM-Leu and CSM-Leu+Naringenin plates. **a-d** Components of naringenin addiction circuits containing promoters $P_{O\text{-}TEF(111)}$, $P_{O\text{-}TEF(136)}$, $P_{O\text{-}LEU2(78)}$, and $P_{O\text{-}GAPDH(88)}$. **e-h** The control set. Growth of *Y. lipolytica* transformants containing promoters $P_{O\text{-}TEF(111)}$, $P_{O\text{-}TEF(136)}$, $P_{O\text{-}LEU2(78)}$, and $P_{O\text{-}GAPDH(88)}$ on CSM-Leu plates. Pictures were taken 3 days after transformation. **i-l** Growth of *Y. lipolytica* transformants containing promoters $P_{O\text{-}TEF(111)}$, $P_{O\text{-}TEF(136)}$, $P_{O\text{-}LEU2(78)}$, and $P_{O\text{-}GAPDH(88)}$ on CSM-Leu+Naringenin plates. Pictures were taken 3 days after transformation.

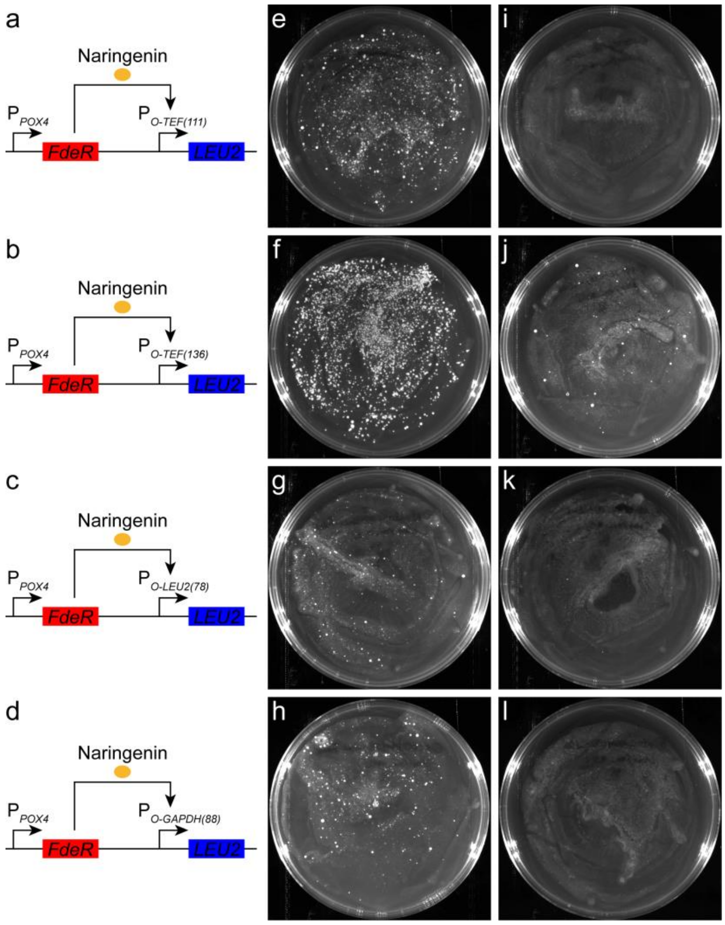


**Supplementary Fig. 3** Analysis of complete and incomplete naringenin addiction circuits in naringenin producing strain NarPro/ASC_Rep. **a-d** Components of naringenin addiction circuits containing promoters $P_{O\text{-}TEF(111)}$, $P_{O\text{-}TEF(136)}$, $P_{O\text{-}LEU2(78)}$, and $P_{O\text{-}GAPDH(88)}$. FdeR was expressed using weaker promoter $P_{POX4}$. **e-h** Growth of NarPro/ASC_Rep containing complete addiction circuits on leucine drop-out CSM-leu plates. *LEU2* expression was controlled by $P_{O\text{-}TEF(111)}$, $P_{O\text{-}TEF(136)}$, $P_{O\text{-}LEU2(78)}$, and $P_{O\text{-}GAPDH(88)}$, respectively. Pictures were taken 3 days after transformation. **i-l** Negative control set. Growth of NarPro/ASC_Rep containing incomplete addiction circuits without FdeR on leucine drop-out CSM-leu plates. *LEU2* expression was controlled by $P_{O\text{-}TEF(111)}$, $P_{O\text{-}TEF(136)}$, $P_{O\text{-}LEU2(78)}$, and $P_{O\text{-}GAPDH(88)}$, respectively. Pictures were taken 3 days after transformation.

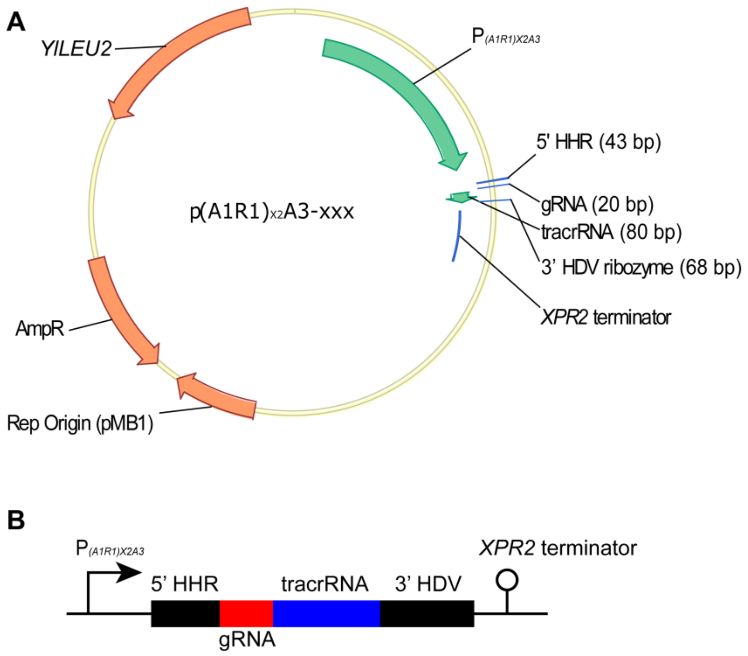


**Supplementary Fig. 4. Genetic structure of plasmid p(A1R1)$_{x2}$A3-xxx.** "xxx" here refers to gRNA. For instance, p(A1R1)$_{x2}$A3-FAS1-2 refers to p(A1R1)$_{x2}$A3 containing gRNA FAS1-2 generating components. **A.** Overall structure of plasmid p(A1R1)$_{x2}$A3-xxx. **B.** Structure of the gRNA generating components. The 5' HHR and 3' HDV ribozyme sites were used for generating mature gRNAs.

### Supplementary tables

#### Supplementary Table 1 gRNA sequences and targeting locations

| gRNA | Sequence (5’-3’) | Targeting locations |
|---|---|---|
| FAS1-1 | ACTCACCAAATCAAATAATG | UTR |
| FAS1-2 | TTTTGCTACAGGAAACAGCG | Intron |
| FAS1-3 | GGTGCAGTTGATGTACAGAG | CDS |
| FAS2-1 | AAACCTGAGCCACAAATCAG | Intron |
| FAS2-2 | GGGCATCACCACCAGCAGCG | CDS |
| FAS2-3 | GAAGGGGACAACAATCAGAG | CDS |
| FabD-1 | GCAGCATTCTTCCCCGGACA | CDS |
| FabD-2 | GAGGCGTTGGATACCACCTG | CDS |
| FabD-3 | CACATGATATACACAAAGCA | Intron |

**Supplementary Table 2 Plasmids used in this paper**

| Plasmid | Annotation |
|---|---|
| pYLXP’ | A frozen stock of our Lab[1]. |
| pYLXP’-Nluc | A frozen stock of our Lab[2]. |
| pYLXP’-dCas9 | A frozen stock of our Lab. |
| pYLXP’-Cre | A frozen stock of our Lab[3]. |
| pYLXP’-URA3-loxP | A frozen stock of our Lab[3]. |
| pΔleu2loxP | Integration at *leu2* site. |
| pΔxpr2loxP | Integration at *xpr2* site. |
| pΔleu2loxP-POX2(1591)-Nluc | Integrating POX2(1591)-Nluc at *leu2* site. |
| pΔleu2loxP-A1R1A3-Nluc | Integrating A1R1A3-Nluc at *leu2* site. |
| pΔleu2loxP-$(A1R1)_{x2}$A3-Nluc | Integrating $(A1R1)_{x2}$A3-Nluc at *leu2* site. |
| pΔleu2loxP-FAS1-1-dCas9 | Integrating gRNA FAS1-1 and dCas9 circuits at *leu2* site. |
| pΔleu2loxP-FAS1-2-dCas9 | Integrating gRNA FAS1-2 and dCas9 circuits at *leu2* site. |
| pΔleu2loxP-FAS1-3-dCas9 | Integrating gRNA FAS1-3 and dCas9 circuits at *leu2* site. |
| pΔleu2loxP-FAS2-1-dCas9 | Integrating gRNA FAS2-1 and dCas9 circuits at *leu2* site. |
| pΔleu2loxP-FAS2-2-dCas9 | Integrating gRNA FAS2-2 and dCas9 circuits at *leu2* site. |
| pΔleu2loxP-FAS2-3-dCas9 | Integrating gRNA FAS2-3 and dCas9 circuits at *leu2* site. |
| pΔleu2loxP-FabD-1-dCas9 | Integrating gRNA FabD-1 and dCas9 circuits at *leu2* site. |
| pΔleu2loxP-FabD-2-dCas9 | Integrating gRNA FabD-2 and dCas9 circuits at *leu2* site. |
| pΔleu2loxP-FabD-3-dCas9 | Integrating gRNA FabD-3 and dCas9 circuits at *leu2* site. |
| pΔleu2loxP-FAS1-2-FAS2-1-dCas9 | Integrating gRNAs FAS1-2 and FAS2-1 and dCas9 circuits at *leu2* site. |
| pΔleu2loxP-FAS1-2-FabD-1-dCas9 | Integrating gRNAs FAS1-2 and FabD-1 and dCas9 circuits at *leu2* site. |
| pΔleu2loxP-FAS2-1-FabD-1-dCas9 | Integrating gRNAs FAS2-1 and FabD-1 and dCas9 circuits at *leu2* site. |
| pΔleu2loxP-FAS1-2-FAS2-1-FabD-1-dCas9 | Integrating gRNAs FAS1-2, FAS2-1, and FabD-1 and dCas9 circuits at *leu2* site. |
| pOTEF(111) | Replacing *TEF* promoter in pYLXP’2 with *OTEF(111)* hybrid promoter. |
| pOTEF(136) | Replacing *TEF* promoter in pYLXP’2 with *OTEF(136)* hybrid promoter. |
| pOLEU2(78) | Replacing *TEF* promoter in pYLXP’2 with *OLEU2(78)* hybrid promoter. |
| pOGAPDH(88) | Replacing *TEF* promoter in pYLXP’2 with *OGAPDH(88)* hybrid promoter. |
| pOTEF(111)-Nluc | Placing *Nluc* under the control of hybrid promoter *OTEF(111)*. |
| pOTEF(136)-Nluc | Placing *Nluc* under the control of hybrid promoter *OTEF(136)*. |

| | |
|---|---|
| pOLEU2(78)-Nluc | Placing *Nluc* under the control of hybrid promoter *OLEU2(78)*. |
| pOGAPDH(88)-Nluc | Placing *Nluc* under the control of hybrid promoter *OGAPDH(88)*. |
| pOTEF(111)-LEU2 | Placing *LEU2* under the control of hybrid promoter *OTEF(111)*. |
| pOTEF(136)-LEU2 | Placing *LEU2* under the control of hybrid promoter *OTEF(136)*. |
| pOLEU2(78)-LEU2 | Placing *LEU2* under the control of hybrid promoter *OLEU2(78)*. |
| pOGAPDH(88)-LEU2 | Placing *LEU2* under the control of hybrid promoter *OGAPDH(88)*. |
| pYLXP'-FdeR | Placing FdeR under the control of promoter *TEF*. |
| pYaliJ1-FdeR | Placing FdeR under the control of promoter *POX4*. |
| pYaliL1-FdeR | Placing FdeR under the control of promoter *IDP2*. |
| pOTEF(111)-FdeR | Placing *FdeR* under the control of hybrid promoter *OTEF(111)*. |
| pOTEF(136)-FdeR | Placing *FdeR* under the control of hybrid promoter *OTEF(136)*. |
| pOLEU2(78)-FdeR | Placing *FdeR* under the control of hybrid promoter *OLEU2(78)*. |
| pOGAPDH(88)-FdeR | Placing *FdeR* under the control of hybrid promoter *OGAPDH(88)*. |
| pOTEF(111)-Nluc-FdeR | pOTEF(111)-Nluc containing *FdeR* under the control of *TEF* promoter. |
| pOTEF(136)-Nluc-FdeR | pOTEF(136)-Nluc containing *FdeR* under the control of *TEF* promoter. |
| pOLEU2(78)-Nluc-FdeR | pOLEU2(78)-Nluc containing *FdeR* under the control of *TEF* promoter. |
| pOGAPDH(88)-Nluc-FdeR | pOGAPDH(88)-Nluc containing *FdeR* under the control of *TEF* promoter. |
| pOTEF(111)-LEU2-FdeR | pOTEF(111)-LEU2 containing *FdeR* under the control of *TEF* promoter. |
| pOTEF(136)-LEU2-FdeR | pOTEF(136)-LEU2 containing *FdeR* under the control of *TEF* promoter. |
| pOLEU2(78)-LEU2-FdeR | pOLEU2(78)-LEU2 containing *FdeR* under the control of *TEF* promoter. |
| pOGAPDH(88)-LEU2-FdeR | pOGAPDH(88)-LEU2 containing *FdeR* under the control of *TEF* promoter. |
| pOTEF(111)-LEU2-POX4-FdeR | pOTEF(111)-LEU2 containing *FdeR* under the control of *POX4* promoter. |
| pOTEF(136)-LEU2-POX4-FdeR | pOTEF(136)-LEU2 containing *FdeR* under the control of *POX4* promoter. |
| pOLEU2(78)-LEU2-POX4-FdeR | pOLEU2(78)-LEU2 containing *FdeR* under the control of *POX4* promoter. |
| pOGAPDH(88)-LEU2-POX4-FdeR | pOGAPDH(88)-LEU2 containing *FdeR* under the control of *POX4* promoter. |
| pOTEF(111)-LEU2-IDP2-FdeR | pOTEF(111)-LEU2 containing *FdeR* under the control of *IDP2* promoter. |
| pOTEF(136)-LEU2-IDP2-FdeR | pOTEF(136)-LEU2 containing *FdeR* under the control of *IDP2* |

| | |
|---|---|
| | promoter. |
| pOLEU2(78)-LEU2-IDP2-FdeR | pOLEU2(78)-LEU2 containing *FdeR* under the control of *IDP2* promoter. |
| pOGAPDH(88)-LEU2-IDP2-FdeR | pOGAPDH(88)-LEU2 containing *FdeR* under the control of *IDP2* promoter. |
| pOTEF(136)-LEU2-OTEF(111)-FdeR | pOTEF(136)-LEU2 containing *FdeR* under the control of *OTEF(111)* promoter. |
| pOTEF(136)-LEU2-OTEF(136)-FdeR | pOTEF(136)-LEU2 containing *FdeR* under the control of *OTEF(136)* promoter. |
| pOTEF(136)-LEU2-OLEU2(78)-FdeR | pOTEF(136)-LEU2 containing *FdeR* under the control of *OLEU2(78)* promoter. |
| pOTEF(136)-LEU2-OGAPDH(88)-FdeR | pOTEF(136)-LEU2 containing *FdeR* under the control of *OGAPDH(88)* promoter. |
| pΔxpr2loxP-OTEF(111)-Nluc-FdeR | Integrating OTEF(111)-Nluc-FdeR at *xpr2* site. Used for testing *OTEF(111)* promoter. |
| pΔxpr2loxP-OTEF(136)-Nluc-FdeR | Integrating OTEF(136)-Nluc-FdeR at *xpr2* site. Used for testing *OTEF(136)* promoter. |
| pΔxpr2loxP-OLEU2(78)-Nluc-FdeR | Integrating OLEU2(78)-Nluc-FdeR at *xpr2* site. Used for testing *OLEU2(78)* promoter. |
| pΔxpr2loxP-OGAPDH(88)-Nluc-FdeR | Integrating OGAPDH(88)-Nluc-FdeR at *xpr2* site. Used for testing *OGAPDH(88)* promoter. |
| pΔxpr2loxP-OTEF(111)-LEU2 | Integrating OTEF(111)-LEU2 at *xpr2* site. Used as control. |
| pΔxpr2loxP-OTEF(136)-LEU2 | Integrating OTEF(136)-LEU2 at *xpr2* site. Used as control. |
| pΔxpr2loxP-OLEU2(78)-LEU2 | Integrating OLEU2(78)-LEU2 at *xpr2* site. Used as control. |
| pΔxpr2loxP-OGAPDH(88)-LEU2 | Integrating OGAPDH(88)-LEU2 at *xpr2* site. Used as control. |
| pΔxpr2loxP-OTEF(111)-LEU2-FdeR | Integrating OTEF(111)-LEU2-FdeR at *xpr2* site. |
| pΔxpr2loxP-OTEF(136)-LEU2-FdeR | Integrating OTEF(136)-LEU2-FdeR at *xpr2* site. |
| pΔxpr2loxP-OLEU2(78)-LEU2-FdeR | Integrating OLEU2(78)-LEU2-FdeR at *xpr2* site. |
| pΔxpr2loxP-OGAPDH(88)-LEU2-FdeR | Integrating OGAPDH(88)-LEU2-FdeR at *xpr2* site. |
| pΔxpr2loxP-OTEF(111)-LEU2-POX4-FdeR | Integrating OTEF(111)-LEU2-POX4-FdeR at *xpr2* site. |
| pΔxpr2loxP-OTEF(136)-LEU2-POX4-FdeR | Integrating OTEF(136)-LEU2-POX4-FdeR at *xpr2* site. |

| | |
|---|---|
| pΔxpr2loxP-OLEU2(78)-LEU2-POX4-FdeR | Integrating OLEU2(78)-LEU2-POX4-FdeR at *xpr2* site. |
| pΔxpr2loxP-OGAPDH(88)-LEU2-POX4-FdeR | Integrating OGAPDH(88)-LEU2-POX4-FdeR at *xpr2* site. |
| pΔxpr2loxP-OTEF(111)-LEU2-IDP2-FdeR | Integrating OTEF(111)-LEU2-IDP2-FdeR at *xpr2* site. |
| pΔxpr2loxP-OTEF(136)-LEU2-IDP2-FdeR | Integrating OTEF(136)-LEU2-IDP2-FdeR at *xpr2* site. |
| pΔxpr2loxP-OLEU2(78)-LEU2-IDP2-FdeR | Integrating OLEU2(78)-LEU2-IDP2-FdeR at *xpr2* site. |
| pΔxpr2loxP-OGAPDH(88)-LEU2-IDP2-FdeR | Integrating OGAPDH(88)-LEU2-IDP2-FdeR at *xpr2* site. |
| pΔxpr2loxP-OTEF(136)-LEU2-OTEF(111)-FdeR | Integrating OTEF(136)-LEU2-OTEF(111)-FdeR at *xpr2* site. |
| pΔxpr2loxP-OTEF(136)-LEU2-OTEF(136)-FdeR | Integrating OTEF(136)-LEU2-OTEF(136)-FdeR at *xpr2* site. |
| pΔxpr2loxP-OTEF(136)-LEU2-OLEU2(78)-FdeR | Integrating OTEF(136)-LEU2-OLEU2(78)-FdeR at *xpr2* site. |
| pΔxpr2loxP-OTEF(136)-LEU2-OGAPDH(88)-FdeR | Integrating OTEF(136)-LEU2-OGAPDH(88)-FdeR at *xpr2* site. |

**Supplementary Table 3 Primers used in this paper**

| Primer | Sequence (5'-3') |
|---|---|
| leu2-up F | ATCCCTAAATTTGATGAAAGCCTAGGGCATAAAATGTGGAGAAGAAATC |
| leu2-up R | CCAACCCGGTCTCTGTCGTCTGTGGATGTGTGTGGTTGTATG |
| leu2-down F | AGCTTTACCGCAGCAGATCCGTCGTTTCTACGACGCATTGATGG |
| leu2-down R | CACTATTGGCCTATGCGGCCGCTGGCACTGAGCTCGTCTAACGG |
| xpr2-up F | ATCCCTAAATTTGATGAAAGCCTAGGGACGAGGACTCGTCCAACGG |
| xpr2-up R | CCAACCCGGTCTCTGTCGTCACCGAGAAGATCAGCGTTCTGTG |
| xpr2-down F | AGCTTTACCGCAGCAGATCCGACGCCAACACCAAGCTGGT |
| xpr2-down R | CACTATTGGCCTATGCGGCCGCGGACTCAGTAATAAGAGCCTCG |
| Nluc F | ACCAGCACTTTTTGCAGTACTAACCGCAGGTCTTCACACTCGAAGATTTCG |
| Nluc R | CATAGCACGCGTGTAGATACTTACGCCAGAATGCGTTCGCAC |
| LEU2 F | ACCAGCACTTTTTGCAGTACTAACCGCAGGAACCCGAAACTAAGAAGACCAAGA |
| LEU2 R | CATAGCACGCGTGTAGATACTTATACACTAGCGGACCCTGCCGGT |
| pPOX2(1591) F | CTAAATTTGATGAAAGCCTAGGGACGACGATATTCCGGTCCCGAAACCC |
| pPOX2(1591) R | CTGCCTCTGAAACTCACCATTCTAGAGGCGTCGTTGCTTGTGTG |
| A3 F | CGAATGGTACGATTCCGCCACAATTGGACATGTTTGTTTTTCCGATC |
| A1R1 R | GATCGGAAAAACAAACATGTCCAATTGTGGCGGAATCGTACCATTCG |
| A1R1_R1 | GGTTTCGGGACCGGAATATCTGGCGGAATCGTACCATTCGC |
| A1R1_F2 | GATATTCCGGTCCCGAAACCCC |
| A1R1_R2 | TGGCGGAATCGTACCATTCGCC |
| gRNA F | TCCCTAAATTTGATGAAAGCCTAGGGACGACGATATTCCGGTCCC |
| gRNA R | CCTTTTATCAGACATAGTCGACTCCTCCGTTATTGTCTCGCTAGC |
| gRNA_FAS1-1 F | GACGAAACGAGTAAGCTCGTCACTCACCAAATCAAATAATGGTTTTAGAGCTAGAAATAG |
| gRNA_FAS1-1 R | ACGAGCTTACTCGTTTCGTCCTCACGGACTCATCAGACTCACCTGCGGTTAGTACTGCAA |
| gRNA_FAS1-2 F | GACGAAACGAGTAAGCTCGTCTTTTGCTACAGGAAACAGCGGTTTTAGAGCTAGAAATAG |
| gRNA_FAS1-2 R | ACGAGCTTACTCGTTTCGTCCTCACGGACTCATCAGTTTTGCCTGCGGTTAGTACTGCAA |
| gRNA_FAS1-3 F | GACGAAACGAGTAAGCTCGTCGGTGCAGTTGATGTACAGAGGTTTTAGAGCTAGAAATAG |
| gRNA_FAS1-3 R | ACGAGCTTACTCGTTTCGTCCTCACGGACTCATCAGGGTGCACTGCGGTTAGTACTGCAA |
| gRNA_FAS2-1 F | GACGAAACGAGTAAGCTCGTCAAACCTGAGCCACAAATCAGGTTTTAGAGCTAGAAATAG |
| gRNA_FAS2-1 R | ACGAGCTTACTCGTTTCGTCCTCACGGACTCATCAGAAACCTCTGCGGTTAGTACTGCAA |
| gRNA_FAS2-2 F | GACGAAACGAGTAAGCTCGTCGGGCATCACCACCAGCAGCGGTTTTAGAGCTAGAAATAG |
| gRNA_FAS2-2 R | ACGAGCTTACTCGTTTCGTCCTCACGGACTCATCAGGGGCATCTGCGGTTAGTACTGCAA |

| | |
|---|---|
| gRNA_FAS2-3 F | GACGAAACGAGTAAGCTCGTCGAAGGGGACAACAATCAGAGGTTTTAGAGCTAGAAATAG |
| gRNA_FAS2-3 R | ACGAGCTTACTCGTTTCGTCCTCACGGACTCATCAGGAAGGGCTGCGGTTAGTACTGCAA |
| gRNA_FabD-1 F | GACGAAACGAGTAAGCTCGTCGCAGCATTCTTCCCCGGACAGTTTTAGAGCTAGAAATAG |
| gRNA_FabD-1 R | ACGAGCTTACTCGTTTCGTCCTCACGGACTCATCAGGCAGCACTGCGGTTAGTACTGCAA |
| gRNA_FabD-2 F | GACGAAACGAGTAAGCTCGTCGAGGCGTTGGATACCACCTGGTTTTAGAGCTAGAAATAG |
| gRNA_FabD-2 R | ACGAGCTTACTCGTTTCGTCCTCACGGACTCATCAGGAGGCGCTGCGGTTAGTACTGCAA |
| gRNA_FabD-3 F | GACGAAACGAGTAAGCTCGTCCACATGATATACACAAAGCAGTTTTAGAGCTAGAAATAG |
| gRNA_FabD-3 R | ACGAGCTTACTCGTTTCGTCCTCACGGACTCATCAGCACATGCTGCGGTTAGTACTGCAA |
| TEF_F2 | GGGTATAAAAGACCACCGTCCCC |
| 26srDNA2s F | ATCCCTAAATTTGATGAAAGCCTAGGCAGACACTGCGTCGCTCCGTCC |
| XPR2_Seq | GGTGTTGGACTCAGTAATAAGAGCC |
| leu2_Inte F | GTGTGCACTCCAACTTTTCACAC |
| xpr2_Inte F | AGCCGTGTTTCGTGACGCAATC |
| leu2_Inte R | GATCATGCACACATAAGGTCC |
| $FAS1_{rt}$ F[1] | GTCTCTGTATGGTCTGTGTCTTG |
| $FAS1_{rt}$ R | GAGTGGAAAGGAGAGGTGATG |
| $FAS2_{rt}$ F | CACTCTCCCTTTTCTCCACATC |
| $FAS2_{rt}$ R | AGCAACCTCAAGACCATCG |
| $FabD_{rt}$ F | GCATCTCAAAGCCTCCAAAC |
| $FabD_{rt}$ R | TGAAATCGGGACGGATCTTG |
| $ACT1_{rt}$ F | GGTATCGTTCTTGACTCTGGTG |
| $ACT1_{rt}$ R | AGGTAGTCGGTAAGATCTCGG |

1. The subscripts "$_{rt}$" refers to primers used for qRT-PCR.
